# Neotropical bats as bioindicators for emerging zoonoses in Central America: A case study identifying *Trypanosoma cruzi* in bats from Belize using metagenomic next-generation sequencing

**DOI:** 10.64898/2025.12.10.693613

**Authors:** Elissa G. Torgerson, Michele Adams, Lauren R. Lock, Molly C. Simonis, Kristin E. Dyer, Amanda Vicente-Santos, M. Brock Fenton, Nancy B. Simmons, Daniel J. Becker, Nicole L. Achee

**Affiliations:** Department of Biological Sciences, College of Science, University of Notre Dame, Notre Dame, Indiana, United States of America; School of Biological Sciences, College of Arts and Sciences, University of Oklahoma, Norman, Oklahoma, United States of America; College of Forestry, Wildlife and Environment, and College of Veterinary Medicine Department of Pathobiology, Auburn University, Auburn, Alabama, United States of America; Department of Biology, Western University, London, Ontario, Canada; Department of Mammalogy, American Museum of Natural History, New York, New York, United States of America; Department of Biological Sciences, Eck Institute for Global Health, University of Notre Dame, Notre Dame, Indiana, United States of America

## Abstract

Emerging zoonoses remain a global public health concern. Surveillance of infectious and vector-borne diseases is vital for predicting and mitigating detrimental effects of zoonotic spillover events. Beyond assessing what microorganisms are circulating in specific environments, it is important to understand how potential reservoir hosts, especially animals such as bats, participate in pathogen transmission. Bats can host and potentially spread infections caused by bacteria, viruses, fungi, and protozoa. However, bats can also act as bioindicators that test positive for pathogenic microorganisms without necessarily contributing to the pathogen replication cycle. Metagenomic next-generation sequencing (mNGS) provides an efficient means to broadly screen for pathogens, although microorganism selectivity can sometimes be lower than targeted approaches. Pairing mNGS results with higher-sensitivity tests such as quantitative PCR (qPCR) can validate results and together these tools provide a relatively fast and reliable method for conducting surveillance. To test this approach, we surveyed the types of microorganisms circulating in Belize by collecting 263 blood samples from 20 different bat species captured in the Orange Walk District in 2019, 2022, and 2023. We used mNGS to initially characterize the microbial communities and qPCR to confirm presence and intensity of human pathogens of interest. We detected 1,430 different microorganisms with some relevance to human or animal health, including the protozoan *Trypanosoma cruzi* which was detected in the phyllostomid bats *Desmodus rotundus* and *Artibeus jamaicensis*. qPCR confirmed the presence and intensity of *Trypanosoma cruzi* in mNGS-positive bat samples. We documented the types of pathogenic microorganisms circulating throughout the bat community in northern Belize to demonstrate the capacity for bats to serve as bioindicators.

**Author Summary:** Tracking the spread of new and emerging zoonotic diseases is a major component of global health research. Pathogen surveillance is a vital part of predicting and reducing the consequences of disease outbreaks. Bats are a diverse group of mammals that can host and potentially transmit many pathogens that pose potential risks to human and environmental health. Our study surveyed blood samples (n=263) from 20 bat species collected from the Orange Walk District of Belize in 2019, 2022, and 2023. Metagenomic next-generation sequencing identified 1,430 different microorganisms that are considered potentially relevant to human or animal health. Among the microorganisms detected was *Trypanosoma cruzi* (*T. cruzi*), the protozoan causative agent of Chagas disease. *T. cruzi* was of particular interest due to its presence throughout the Americas and relevance to public health. We surveyed the types of microorganisms circulating throughout bat populations in northern Belize to demonstrate the ability of bats to act as bioindicators.

## Introduction

Risks associated with emerging zoonoses are increasing as populations of humans and livestock grow and the world continues to become more interconnected and interdependent. Instability of various kinds (political, economic, environmental) encourages movements of people, while limited medical resources jeopardize community health, and climate change influences vector distributions [1]. Increasing atmospheric temperatures are limiting some resources and decreasing habitable land [2]. For these reasons, the expansion of disease surveillance and neglected disease research are a necessity, as risk of infection is no longer isolated to contained regions. Human mobility inevitably increases pathogen mobility [3]. Infection rates are likely to rise as proximity between human populations declines and vectors acclimate to new environments. In addition, increases in unusual and severe weather conditions can facilitate the spread of pathogens through contaminated resources (i.e., water, food, air, and livestock) [4,5]. Changes in land use have also facilitated greater overlap of people with wildlife and concomitantly led to increases in zoonotic spillover risk [6–8]. Humans are not the only ones being affected by growing pressures resulting from climate change and habitat conversion, as many animal and non-animal organisms, such as bats, are having to adapt to survive [9]. Bats are important animals to consider when conducting surveillance because they can host a large number of pathogens, are known to participate in cross-species pathogen transmission, and they are reservoirs for important zoonotic pathogens like rabies lyssavirus, Hendra and Nipah henipaviruses, and Marburg filovirus [10–12].

Bats are a diverse and wide-spread group of mammals that inhabit a variety of ecosystems and exploit both urban and non-urban settings [13]. Healthy bat populations are essential for maintaining balanced ecosystems as they provide services like seed dispersal, plant pollination, and agricultural pest predation [7,14,15]. By participating in activities that support ecosystem biodiversity, bats contribute positively to the socio-economic status of the regions they inhabit [16]. While bats provide many ecological benefits, close proximity to humans increases the likelihood of a zoonotic spillover event [7,17]. Climate change, the destruction of natural habitat, and rising competition for natural resources have played notable roles in altering bat behavior [18]. As a result, it is becoming increasingly common for bats to reside in agricultural and urban environments and alter their diet to accommodate their change in habitat [19,20].

The increase in bats residing in urban and agricultural environments can be detrimental to bat health and, in turn, potentially human and domestic animal health [7,21–23]. For example, as the diets of bats shift to accommodate new food sources, their microbiota composition also changes, often with gut biodiversity declining in anthropogenic environments [24–26]. For some bat species, moving into urban and agricultural environments is also associated with a decline in immunity, potentially increasing susceptibility to infections and extending infectious periods [27,28]. Foraging within anthropogenic environments has especially affected the immune profiles of common vampire bats (*Desmodus rotundus*) because of their exclusively blood diet. While the effects of this behavioral shift are not entirely understood, it has been observed that when vampire bats switch to primarily consuming the blood of livestock, their immune systems can switch to favoring investment in innate immunity, as opposed to adaptive immunity [29].

Robust adaptive immune responses are crucial for host defense against viral pathogens, which means an increase in the reliance of bats on domestic animal prey could compromise their ability to fight-off infection [20,29,30]. Greater reliance on more abundant and stable anthropogenic food sources at the cost of lowering overall immunity has been observed across many bat species. For example, Australian flying foxes (*Pteropus* spp.) rely on poorer quality fruit and nectar resources in urban and agricultural habitats in response to food shortage events, leading to higher stress, decreased immunity, and amplified pulses of Hendra virus shedding and spillover to horses [31–33]. In addition to modifying bat immunity, foraging in urban and agricultural environments also increases overlap among bats, domestic animals, and humans, increasing opportunities for pathogen spillover.

While bats may not necessarily have a greater fraction of zoonotic viruses than other mammal or avian taxa [34], bats can serve as competent hosts for some virulent human-infecting pathogens, possess high antibody gene diversity, and have remarkable immune tolerance compared to other mammals [35–37]. The combination of these immunological factors and propensity to forage and even roost in urban and agricultural environments fosters opportune conditions for zoonotic spillover [38,39]. As such, it is vital that locations and situations that place humans at higher risk of zoonotic spillover (i.e., tropical climate, poor sanitation, close human to animal proximity, natural disasters) are recognized, people are informed of potential threats, and steps are taken to minimize risk [7]. When planning to prevent potential zoonotic spillover, vital information includes understanding the diversity of pathogens present in both wildlife hosts and the environment. Here we address this concern by reporting results of a comprehensive survey of potentially pathogenic microorganisms detected in bat blood samples collected in Belize.

Current surveillance methods used both in and outside of Belize primarily employ targeted approaches such as PCR [40–42], with downstream applications including risk maps and sequencing samples collected from vectors found within the vicinity of confirmed human cases of specific diseases [43,44]. While such approaches are helpful for reliably detecting specific microorganisms of interest, they lack the ability to provide a comprehensive overview of the microorganisms that a host could potentially transmit. Metagenomic next-generation sequencing (mNGS) can detect low concentrations of total nucleic acids (TNA) from a wide variety of microorganisms [45–47], yet it is an underutilized tool for surveillance research.

Here we used mNGS to profile TNA of blood samples from a diverse range of bat species in Belize, including some that have close interactions with humans and domestic animals (e.g., *Desmodus rotundus*). We referenced the National Center for Biotechnology Information [48] and the Chan Zuckerberg Identification Database [49] to identify human and animal pathogens, which we confirmed using qPCR. We were especially interested in attempting to detect *Trypanosoma cruzi*, the causative agent of Chagas disease, given its relevance to human health in Belize and throughout the Americas [44,50,51]. While *T. cruzi* is transmitted to humans by triatomine vectors, bats have been implicated in the Chagas disease cycle throughout the Americas [52–55]. Currently, the primary non-invasive surveillance method for detecting Chagas disease in Belize is through community collections of *Triatoma dimidiata* sensu lato (Reduviidae: Triatominae), which is the predominant Chagas disease vector in the country [40,44]. Community collections rely on residents of specific study areas capturing insects that resemble *Triatoma dimidiata* sensu lato and storing them until the Belize Ministry of Health can collect, verify the identity, and test each insect for *T. cruzi*. Recently, we used targeted PCR to show that *T. cruzi* circulates in *Desmodus rotundus* in Belize, such that other sympatric bat species may also play a role in local transmission cycles [56]. With this in mind, targeting bats as bioindicators could offer an efficient, secondary, non-invasive method for Chagas disease surveillance in Belize.

## Methods

### Bat sampling

During two weeks in April–May 2019, 2022, and 2023, we sampled bats within the Lamanai Archeological Reserve, Orange Walk District, Belize, as part of broader studies on bat immunology and infectious disease [21,57,58]. As described previously, we used mist nets and harp traps to capture bats along flight paths and occasionally at the exits of roosts, typically from 19:00 hours until 24:00 hours. Bats were kept in clean individual cloth bags prior to processing and were identified and sexed based on morphology [59]. We collected small (<1% body mass) volumes of blood by lancing the propatagial vein with sterile needles (26–30G) followed by collection with heparinized capillary tubes. Bats were released following processing. For this study, blood was stored on Whatman FTA cards at room temperature until –20°C storage at the University of Oklahoma. Field procedures were performed according to guidelines for the safe and humane handling of bats published by of the American Society of Mammalogists [60] and were approved by the Institutional Animal Care and Use Committees of the American Museum of Natural History (AMNHIACUC-20190129 and -20210614) and the University of Oklahoma (2022–0197). Fieldwork and sampling were authorized by the Belize Forest Department under permits FD/WL/1/19(06), FD/WL/1/19(09), FD/WL/1/21(12), and FD/WL/1/22(54). For this study, we analyzed 263 blood samples from 20 different bat species in the Belize study region (Fig 1).

**Fig 1.**
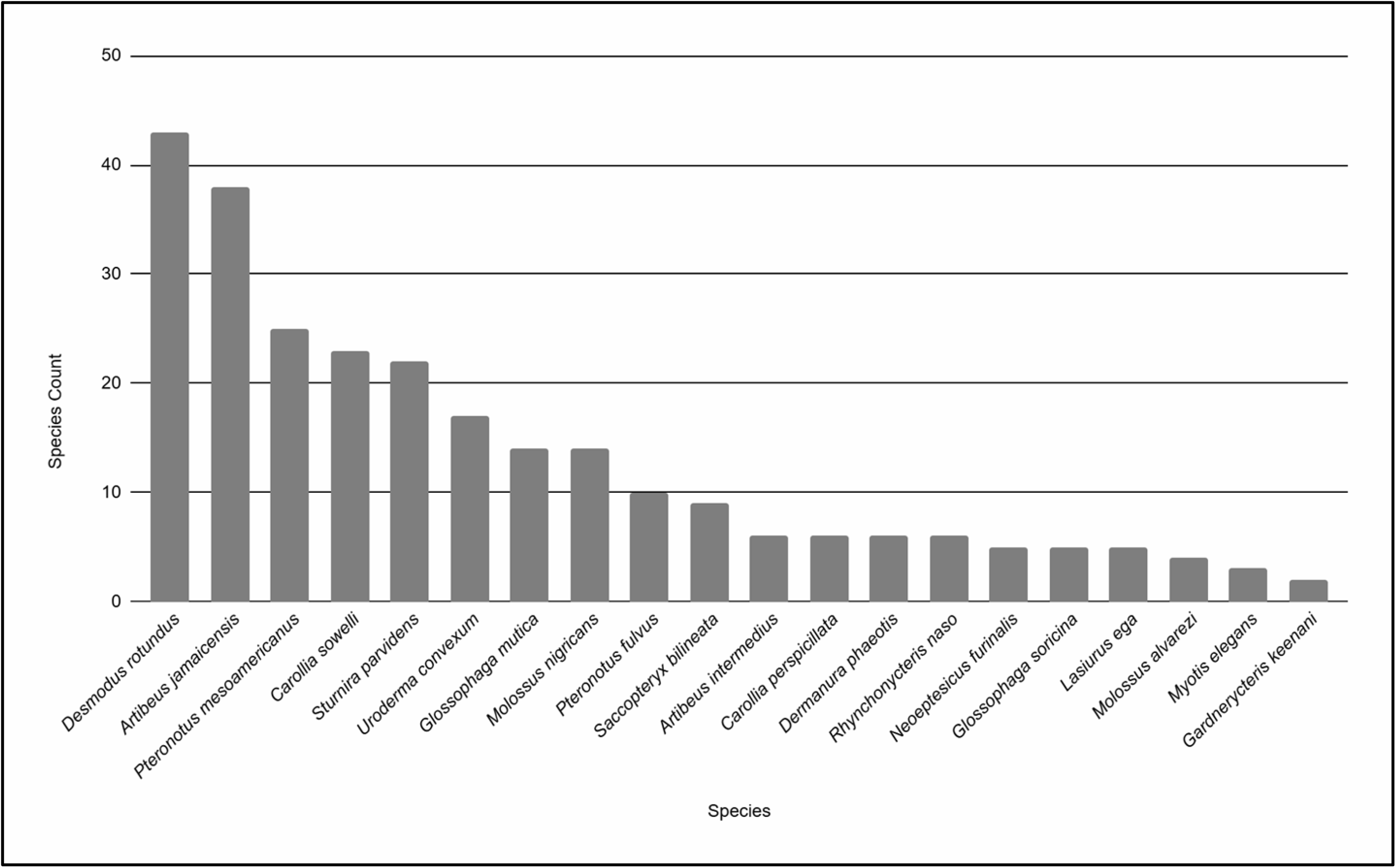
Species of bats sampled. Total number of individual bat species collected in Lamanai, Belize from which blood samples were tested using metagenomic next-generation sequencing (mNGS). Refer to Supporting Files Table 1 for a list of the numbers of each bat species collected in Belize.

We processed and tested all samples according to procedures written by the Remote Emerging Disease Intelligence-NETwork [61–63]. Some changes were made to operating procedures between the years 2023 and 2024, these are noted below.

### TNA extraction

We extracted and sequenced samples over the course of a two-year period. Bat samples collected in 2019 were tested in 2023, while samples collected in 2022 and 2023 were tested in 2024. For all samples, we punched three 3-mm sections containing samples of dried blood spots from FTA cards into a 1.5 mL microcontainer containing zirconium oxide beads. Nucleic acid extraction began by adding 20 µL of proteinase K from an Indical IndiMag Pathogen Kit (Indical BioScience, Florida, USA). For the samples processed in 2023, 500 µL of Qiagen QIAcard FTA Wash buffer (Qiagen, Hilden, Germany) was added to each sample. For samples processed in 2024, 200 µL of Qiagen ATL-DX buffer, and 200 µL of Qiagen QIAcard FTA Wash buffer was added to each of the samples. Samples were vortexed and incubated for 1 hour at 56°C and 1400 rpm. A ThermoFisher KingFisher Flex System (ThermoFisher Scientific, Massachusetts, USA) and an IndiMag Pathogen kit were used for extraction. DNA and RNA concentration measurements were recorded and 75 µL of TNA from each sample was stored for future use.

### Complementary DNA synthesis

Only samples tested in 2024 were used for complementary DNA (cDNA) synthesis. cDNA synthesis was performed using 8 µL of extracted TNA. Invitrogen ezDNase™ Enzyme (ThermoFisher Scientific, Massachusetts, USA) was added to the total nucleic acid, digesting the DNA and allowing the RNA to remain intact. Next, NEBNext^®^ Ultra™ II RNA First Strand Synthesis Module (New England Biolabs (NEB), Massachusetts, USA) and NEBNext^®^ Ultra™ II Non-Directional RNA Second Strand Synthesis Module enzymes were added to create double-stranded cDNA which was then purified. Ten (10) µL of cDNA was combined with 10 µL of genomic DNA (gDNA) from its corresponding sample, producing a total volume of 20 µL for library preparation.

### gDNA and TNA sequencing

For sequencing library preparation, we used 20 µL of gDNA for the samples tested in 2023 and 20 µL of TNA for the samples tested in 2024. Fragmented DNA was subjected to end-prep repair using NEBNext^®^ Ultra™ II End Repair/dA-Tailing Module and NEBNext^®^ FFPE DNA Repair Mix. Barcodes were added from a Native Barcoding Kit 96 V14 (Oxford Nanopore Technologies (ONT), Oxford, UK). After samples were purified, a library pool for multiplex sequencing was created by combining 12 µL from each of the barcoded samples. Adapter ligation was performed using Native Adapter from the ONT Native Barcoding Kit combined with ligase and ligation reaction buffer from the NEB Quick Ligation™ Kit. After washing the library sample using ONT Short Fragment Buffer, 13 µL of eluate was obtained using ONT Elution Buffer. Library concentration was then measured using 1 µL of sample and a Qubit 4 Fluorometer (ThermoFisher Scientific, Massachusetts, USA).

A mNGS approach was used to test samples. Samples were sequenced on either a ONT GridION or PromethION P2 Solo instrument. Both the GridION and the PromethION used MinKNOW operating software v21.11.7 for high accuracy basecalling. Sequencing ran for 48 hours, and the minimum read length was 200 bp. Sequencing data was then pushed through the REDI-NET Custom Pipeline, which uses a nucleotide-based read classifier to identify pathogens of interest to the genus or species level. At the time of sample sequencing, the REDI-NET Pipeline used EDGE v2.4.1 for barcode trimming, removing reads with q-scores less than 9, and filtering out host DNA [64]. DIAMOND v2.0.14 was used to align reads to the National Center for Biotechnology Information (NCBI) Reference Sequence (RefSeq) database, and MEGAN6 v6.23.2 was used to visualize data and record taxon read counts [65]. Identifications of all microorganisms were confirmed using Blast+ v2.13.0 [66]. For samples of interest that tested positive using mNGS, such as *T. cruzi* (see *Results*), pathogen confirmation was completed using qPCR.

### qPCR confirmation

We used qPCR to confirm any mNGS-positive samples for *T. cruzi*, given the medical importance of this pathogen [44,50]. The PCR mix for *T. cruzi* contained 1x TaqMan™ Fast Advanced Master Mix (ThermoFisher Scientific, Massachusetts, USA), 0.75 μM of primer 1 (5’ AGT CGG CTG ATC GTT TTC GA 3’), primer 2 (5’ AAT TCC TCC AAG CAG CGG ATA 3’), TaqMan™ probe (5’ 6-FAM-CAC ACA CTG GAC ACC AA-QSY 3’) [67], 4 μL template DNA, and nuclease-free water, totaling a volume of 20 μL. Both positive and negative controls were run with samples. Negative controls were nuclease-free water, and template DNA samples that when sequenced tested positive for a *Trypanosoma* species, but negative for *T. cruzi*. The positive control used was gDNA extracted from the *T. cruzi* parasite. To complete *T. cruzi* extraction, a total of 80 µL of ATL-DX Lysis Buffer was transferred into a pre-labeled 1.5 mL bead tube containing 2 scoops of 0.1 mm zirconium oxide beads. A total of 320 µL of the *T. cruzi* parasite was added to the tube and vortexed on its side for 1 minute. The tube was put on ice and allowed to settle for 1 minute before being vortexed on its side again for another minute. The sample was centrifuged at 120 x g for 5 minutes and 350 µL of supernatant was transferred to a new microcentrifuge tube, avoiding bead carryover. Extractions were completed for 1:100 and 1:1,000 *T. cruzi* sample dilutions. DNA and RNA concentrations of positive controls were recorded and gel electrophoresis confirmed positive control efficacy. Samples were tested on a ThermoFisher QuantStudio™ 7 Flex programmed to run at 95°C for 15 minutes, then cycle 45 times at 95°C for 10 seconds, and 54°C for 1 minute. Fluorescence was measured at the end of each cycle at 54°C. Fragments were approximately 166 base pairs [68].

## Results

Our mNGS pipeline detected a total of 1,430 different microorganisms from 10,878 sequencing reads across the 20 sampled Belize bat species. Of the 1,430 different microorganism species, 124 (8.67%) were identified as having some relevance to public health because of their current or potential ability to cause human infection [48,49]. Figure 2 depicts the percentage of the microorganisms identified as having the potential to affect human health.

**Fig 2.**
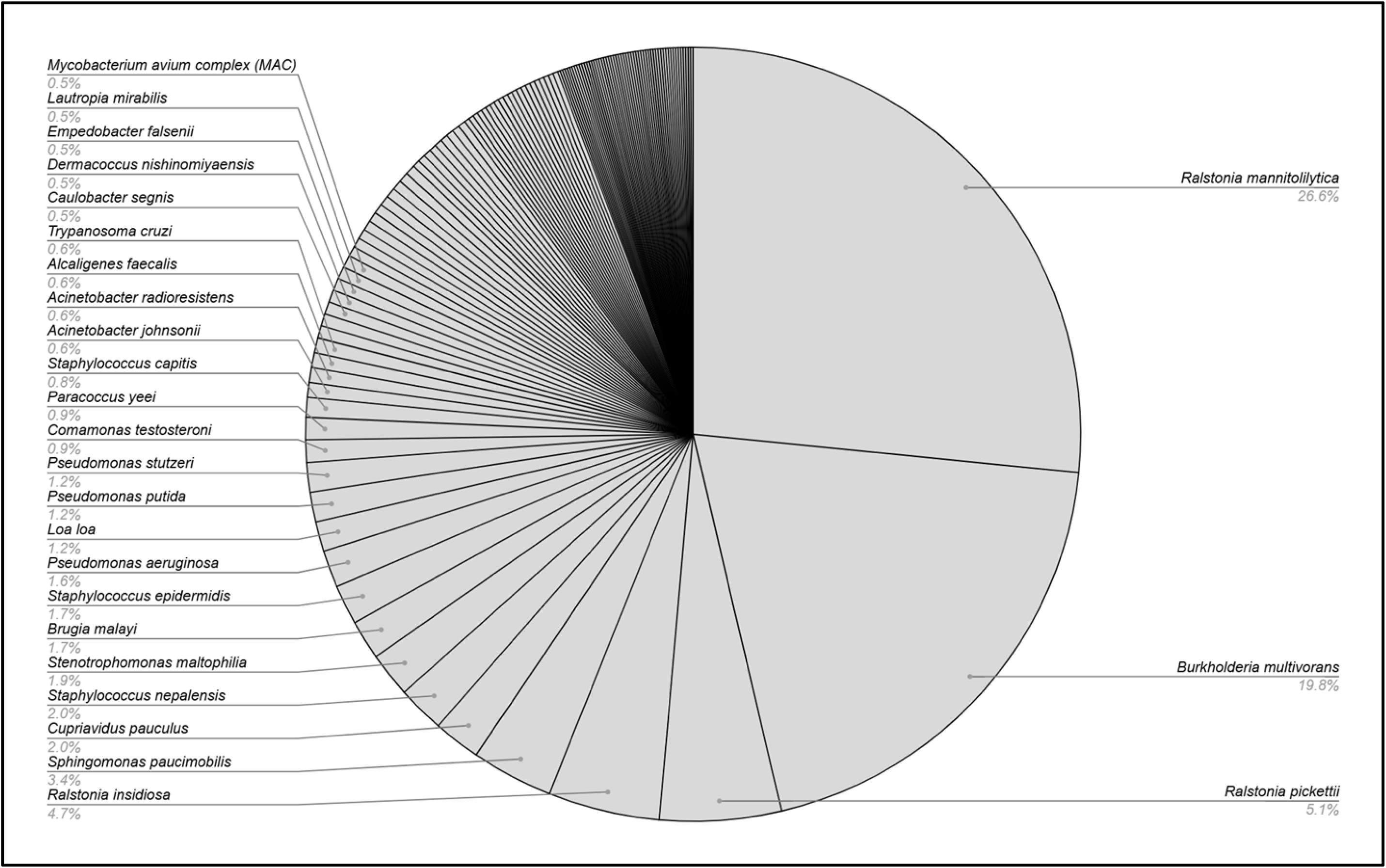
**Pathogens detected in bat blood samples.** The percentage of each pathogen group identified in bat blood samples (n=263) as posing potential risk to human health. Refer to Supporting Files Table 2 for a full microorganism list including pathogens detected in <0.5% of samples. Refer to Supporting Files Figure 1 to view microorganisms present in at least 10% of blood samples.

The most abundant microorganism with the greatest distribution was *Ralstonia mannitolilytica*, appearing in 97.7% of samples in our study. *T. cruzi* was detected in six bat samples across multiple species including *Desmodus rotundus*, *Neoeptesicus furinalis*, *Artibeus jamaicensis*, and *Sturnira parvidens*. The *Trypanosoma* genus was also detected in an additional five bat samples. Of the six samples that tested positive for *T. cruzi* using mNGS, only four tested positive using qPCR (Fig 3).

**Fig 3.**
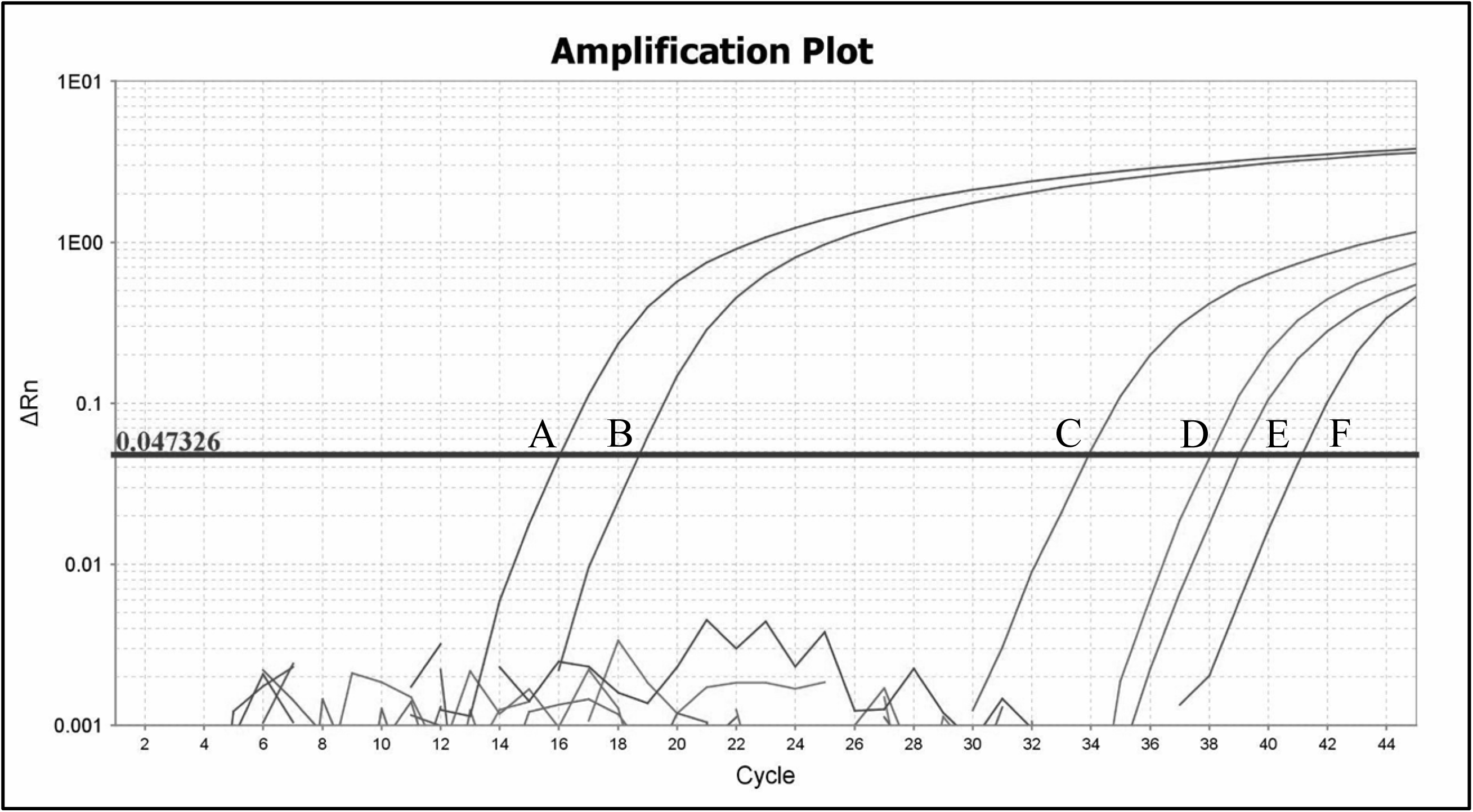
***Trypanosoma cruzi* qPCR amplification plot.** (A) 1:100 C+, positive control, 2 μL 1:100 dilution *Trypanosoma cruzi* gDNA; (B) 1:1,000 C+, positive control, 2 μL 1:1,000 dilution *Trypanosoma cruzi* gDNA; (C) OBZB-00154, 2.0 μL DNA from sample positive for *Trypanosoma cruzi* via mNGS; (D) OBZB-00254, 2.0 μL DNA from sample positive for *Trypanosoma cruzi* via mNGS; (E) OBZB-00161, 2.0 μL DNA from sample positive for *Trypanosoma cruzi* via mNGS; (F) OBZB-00095, 2.0 μL DNA from sample positive for *Trypanosoma cruzi* via mNGS. Refer to Supporting Files Table 3 for a full list of qPCR amplification plot results.

## Discussion

Emerging zoonoses are global public health concerns exacerbated by growing pressures from climate and land use change. Surveillance of infectious and vector-borne diseases is essential for mitigating the detrimental effects of zoonotic spillover. It is important to understand not only which microorganisms are circulating the environment, but also which hosts are (or could become) reservoirs of these infections. Bats can spread an array of infections caused by bacteria, viruses, fungi, and protozoa, although it is not fully understood the extent to which bats participate in zoonotic spillover [24,34]. The primary goal of this study was to determine what microorganisms are circulating throughout bat communities residing in the Lamanai Archeological Reserve located in northern Belize. mNGS was the preferred method for conducting this surveillance project, given that it can broadly screen for an array of microorganisms without the use of specific primers. In total, 1,430 different microorganisms considered potentially relevant to the health of humans and/or animals were detected using this shotgun approach. One pathogen of particular interest that we detected was *T. cruzi*, which was confirmed in four out of six samples using qPCR. Surveying bats via mNGS provides broad information on both target and non-target pathogens, providing a cost-effective method of monitoring infectious disease risks in a diverse and abundant mammal group. In addition to serving as sentinels of possible zoonotic risks, sampling bats for *T. cruzi* could provide a valuable alternative to direct sampling of triatomines, given uncertainty in whether bats could directly transmit this pathogen to humans. Monitoring work will need to weigh the costs and benefits of bat versus triatomine sampling for this specific pathogen [40].

The presence of microorganisms relevant to humans, even when direct transmission is unlikely, raises questions about what types of bat and pathogen dynamics are at play. While *T. cruzi* has several routes of transmission, the primary one occurs when the feces or urine of triatomine bugs come into contact with host mucous membranes or a region of the host body where the skin barrier has been broken by a bug bite [69]. Infection can also occur through blood-based and vertical transmission [70,71]. While there is molecular evidence that supports *T. cruzi* having evolved from a bat trypanosome precursor, sufficient evidence does not exist to demarcate bats as reservoirs for *T. cruzi* [72]. However, some findings do suggest that infected bats could inadvertently transmit *T. cruzi* to other animals, even if bats are the dominant source of pathogen maintenance in the host community. For example, there have been instances of *T. cruzi* spreading via congenital transmission [73] and by oral transmission [74]. Related to this latter transmission route, *T. cruzi* has been detected in the saliva of vampire bats and other sympatric frugivorous bat species [53] which could be the result of food resources contaminated by *T. cruzi* parasites [54]. However, the detection of microorganisms in only blood samples here raises questions about how and why these pathogens are circulating within bat communities. Bats have a distinct tolerance of some pathogens compared to other mammals [36,75], although current understandings of how bat immune systems function and respond to infection has been skewed towards host-viral interactions. Limited work has addressed the bat immune response to protozoan infections, although recent work on vampire bats has suggested bats have at least some level of immune tolerance against *T. cruzi* [56].

Previous reports that have detected *T. cruzi* in bat populations across the Americas suggest diet and behavior could contribute to the likelihood of bats testing positive for this infection. One study from Colombia detected higher prevalence of *T. cruzi* discrete typing units in cave-dwelling bats compared to other classes of discrete typing units [76], while a study from Brazil found that fruit bats were more likely to test positive for *T. cruzi* compared to bats from other dietary guilds [77]. In Ecuador, nectivorous/omnivorous *Glossophaga soricina* residing in a peri-urban environment tested positive for *T. cruzi* [78], while a study from Mexico detected *T. cruzi* in two frugivorous bat species: *Artibeus jamaicensis* and *Sturnira parvidens* [79]. Among these studies, *T. cruzi* prevalence was relatively low, ranging between 1% and 17% of individuals. Comparatively, our results fit within this range, as we confirmed the presence of *T. cruzi* in ∼1.5% of bats sampled at our site. Most of these positives were from vampires (*Desmodus rotundus*), with one positive originating from the frugivore *Artibeus jamaicensis*.

While the factors that increase or decrease *T. cruzi* infection risk in bats remain poorly understood, detection of this pathogen in blood- and fruit-eating bat species provide further support for the role that foraging ecology might participate in bat–trypanosome dynamics. However, more research is required to determine specifically what factors contribute to the perpetuation of *T. cruzi* in bat communities and the environment.

When considering how and why bat blood samples are testing positive for *T. cruzi*, it is important to examine the issue from both ecological and methodological standpoints. mNGS is a relatively new technology, and it is possible that errors could have occurred with sequencing [80]. mNGS is good at detecting multiple pathogens at once without the need for primers [47], and it is also able to identify pathogens that are typically considered more difficult to detect by conventional pathogen identification methods [45,81]. However, there is no universal bioinformatic pipeline used to process mNGS sequencing data, which can cause uncertainty when interpreting results as they can vary across laboratories or between sequencing runs [82]. The increased sensitivity and broad, unbiased approach of mNGS can also increase the chance of false positives [83,84]. Sequencing misalignments and environmental contamination can also make differentiating between pathogenic and non-pathogenic microorganisms (especially those that are genetically related) difficult when interpreting results. In addition, mNGS is typically slower than target approaches like PCR when identifying antimicrobial-resistant genes and mNGS is generally more expensive compared to other pathogen detection methods [85]. However, when used in conjunction with target approaches like PCR, our study has found mNGS to be a reliable and efficient method for pathogen identification.

After confirming the presence of *T. cruzi* in our bat samples, we returned to the question of how and why this pathogen is circulating in bat communities. Integrating information about host-pathogen dynamics could be crucial for understanding how pathogenic microorganisms can be detected within the blood of hosts in cases where the host does not become ill and pathogen replication does not take place. One idea is that hosts are acting as bioindicators and not reservoirs. Bioindicators are generally defined as biotic or abiotic sources that respond to changes in their environment [86], while disease reservoirs are animate or inanimate sources that typically harbor disease-causing organisms and serve as potential sources of disease outbreaks [87]. We are defining bats as bioindicators as opposed to reservoirs because bats are unlikely to cause a Chagas disease outbreak, yet *T. cruzi* can still be detected in their blood indicating if levels circulating in the environment have changed. Overall, this study has demonstrated the capacity for bats to be bioindicators for Chagas disease, while also displaying how shotgun approaches like mNGS can be utilized in combination with targeted approaches like PCR to confirm the presence of certain pathogens of interest. Distinguishing whether host species are functioning as bioindicators or reservoirs is vital when conducting surveillance research because it provides information that can be used to evaluate public health risk and design effective mitigation strategies.

## Acknowledgements

We thank Mark Howells, Karen Gonzalez, Neil Duncan, and staff of the Lamanai Field Research Center for assistance with field logistics and permits. We also thank the many colleagues who helped capture bats during 2019, 2022, and 2023 bat research trips in Belize. We also thank Bret Demory for assistance with sample preparation. Finally, we would like to thank the REDI-NET Consortium for developing the technology and standard operating procedures used for sample processing, testing, and data analysis [https://redi-net.nd.edu].

## Funding

This project was funded by the National Geographic Society (NGS-55503R-19), Research Corporation for Science Advancement (RCSA; Subaward No. 28365, part of a USDA Non-Assistance Cooperative Agreement with RCSA Federal Award No. 58-3022-0-005), National Science Foundation (DBI 2515340), and the Remote Emerging Disease Intelligence-NETwork (REDI-NET). This work is supported by the US Army Medical Research and Development Command under Contract No. W81XWH-21-C-0001, W81XWH-22-C-0093, HT9425-23-C-0059, and HT9425-24-C-0072. The views, opinions, and/or findings presented here are those of the authors and should not be taken to reflect the official policy or position of the U.S. Department of the Navy, U.S. Department of the Army, U.S. Department of Defense, nor that of the U.S. Government. DJB was also supported by the Edward Mallinckrodt, Jr. Foundation.

## Competing interests

Authors have declared no competing interests.

## Supporting information

**S1 Fig.**
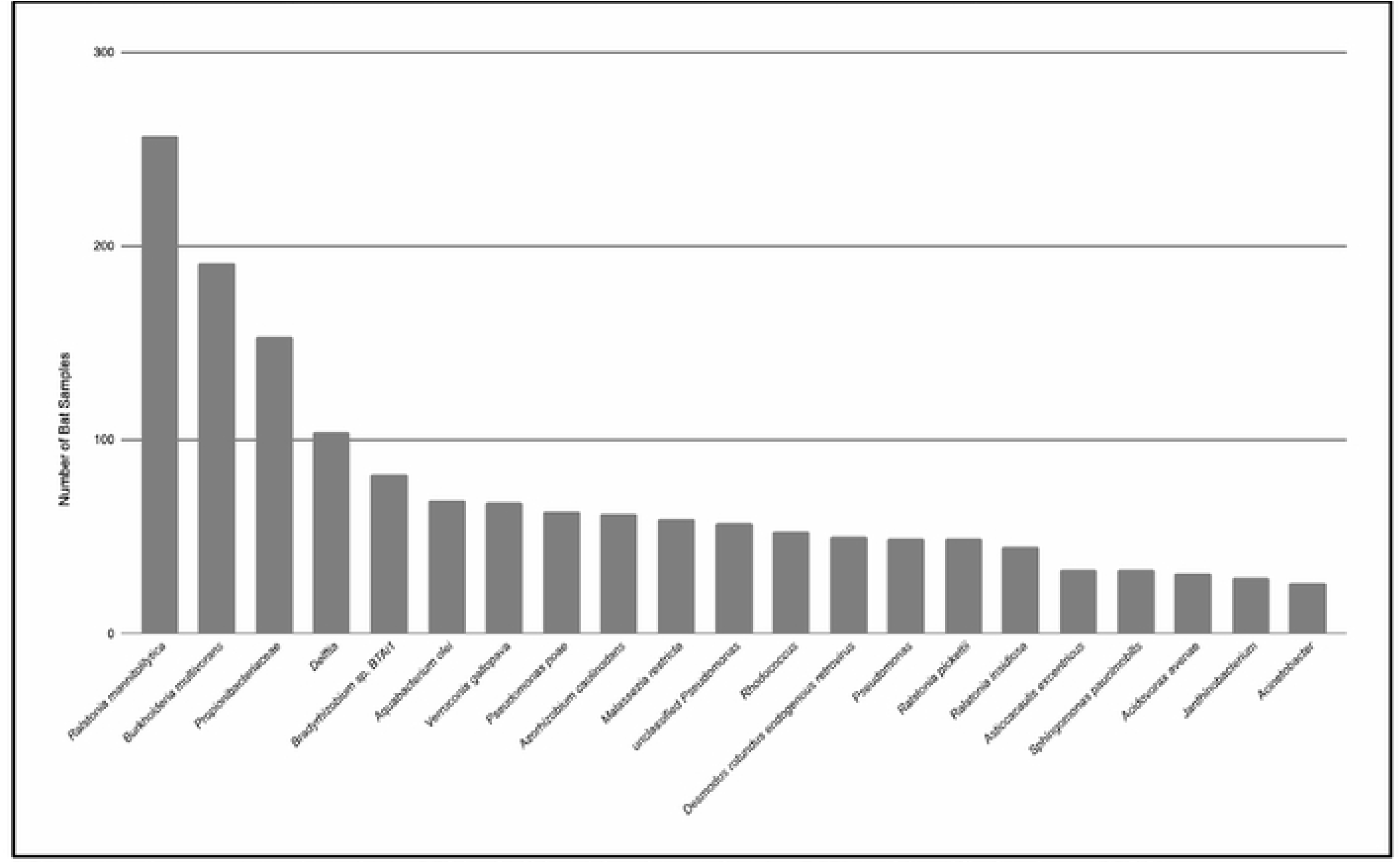
Microorganisms detected in at least 10% of bat blood samples. Microorganisms present in at least 10% of all bat blood samples (n=263) as identified using mNGS.

**S1 Table. Bat species collected in Belize, Central America from which samples were tested using metagenomic next-generation sequencing.**

**S2 Table. Microorganisms detected in blood samples using metagenomic next-generation sequencing.**

**S3 Table. Quantitative PCR amplification plot results for controls and samples that tested positive for Trypanosoma cruzi using metagenomic next-generation sequencing.** C- water, no template control; 1:100 C+, positive control; 1:1,000 C+, positive control; OBZB-00171, negative control; OBZB-00095; OBZB-00121; OBZB-00154; OBZB-00161; OBZB-00165; OBZB-00254.

## Notes

### Competing Interest Statement

The authors have declared no competing interest.

